# Structural selection is revealed only at meso‑structural scales

**DOI:** 10.64898/2026.03.31.715597

**Authors:** Adán Miranda-Pérez, Citlalli H. Mendoza Reyes

## Abstract

Ecological networks are often analyzed as aggregated structures, an approach that has yielded important insights but implicitly assumes that selection acts uniformly across communities. We refine this perspective by showing that structural selection becomes detectable only when analyses focus on meso-structural scales. Using a detailed trophic network, we quantified local structural environments through order-2 egonets and evaluated how structural traits shape interaction geometry. Aggregated representations captured broad patterns but showed no structural gradients, whereas egonets revealed strong axes of differentiation. Hierarchical asymmetry emerged as the dominant meso-structural trait, with local connectivity contributing secondary structure. Principal component analyses and Manhattan distances showed that meso-structural domains retain the heterogeneity through which selection acts. Structural selection was sparse but concentrated in hierarchical domains, identifying the meso-structural scale as the level at which evolutionary signals become detectable in ecological networks. Together, these results position structural selection as an evolutionary process acting on ecological structure, expanding how selection can be conceptualized in complex systems.

**One-Sentence Summary:** Structural selection becomes detectable only at meso-structural scales, revealing evolutionary gradients that vanish in aggregated networks and expanding how selection is conceptualized in structured ecological systems.

**Teaser text:** Ecological networks are usually analyzed as large, aggregated structures, but this perspective hides the evolutionary signals that operate at finer scales. By zooming into meso-structural domains, local neighborhoods captured through order-2 egonets, we uncover strong structural gradients that disappear in whole-network representations. These meso domains reveal where structural selection operates, exposing hierarchical asymmetry as the dominant axis of differentiation. Our results show that aggregated networks smooth out the heterogeneity through which selection acts, whereas meso-structural environments retain the variation that shapes interaction geometry. This work reframes how we detect evolutionary processes in ecological networks and shows that structural selection is sparse, localized, and fundamentally meso-structural, converging with classical natural selection when selection is mediated by interaction geometry—a perspective consistent with recent theoretical work on enemy-tracking constraints in tri-trophic systems by Miranda-Pérez et al. in prep.

## Introduction

Ecological networks are widely used to describe the structure of trophic interactions and to infer the processes that shape community dynamics (Bascompte & Jordano, 2007; Ings et al., 2009; Pascual & Dunne, 2006). These networks are almost always analyzed in aggregated form, collapsing all interactions into a single global topology. This practice assumes that the structural environment experienced by each species is adequately represented by whole-network metrics such as trophic level, centrality, or modularity (Allesina & Tang, 2012; Stouffer et al., 2005; Williams & Martinez, 2000). Yet ecological interactions are inherently local: species encounter only a subset of potential partners, and the structural constraints that shape these encounters operate at scales far smaller than the full network (Thompson, 2005; Tylianakis et al., 2010). This mismatch between analytical scale and ecological reality has important consequences. Aggregated networks often appear homogeneous, with weak or inconsistent structural gradients (Bartomeus et al., 2016; Poisot, 2016). As a result, attempts to detect selective pressures acting on network-embedded traits (such as hierarchical position, asymmetry, or local centrality) frequently yield ambiguous or null results (Gómez et al., 2010; Guimarães et al., 2017; Johnson et al., 2014). Rather than indicating that structural selection is weak or absent, these outcomes may reflect the fact that aggregation can mask the variation through which selection acts. An alternative possibility, therefore, is that the relevant structural heterogeneity is being obscured by analytical scale.

We propose that structural selection emerges at meso structural scales: intermediate interaction neighborhoods that retain the heterogeneity required for selective gradients to form. We define the meso-structural scale as the local interaction environment surrounding each species, captured through order-2 egonets that include both direct and indirect trophic partners. Although earlier studies identified local structural patterns (Cosmo et al., 2023; Olesen et al., 2010; Saavedra & Stouffer, 2013; Stouffer et al., 2012), they did not formalize a meso-structural scale nor evaluate whether selection operates specifically on variation at this level. Egonets preserve the structural context in which interactions occur while avoiding the information loss inherent in whole-network projections. Importantly, this framework reveals a conceptual convergence between structural selection and classical natural selection: both processes act by filtering variation through differential persistence. In ecological networks, meso structural domains provide the structural variation upon which selection can act, analogous to heritable traits in organismal evolution. This parallel clarifies how structural selection complements, rather than replaces, organism level natural selection(Miranda-Pérez et al., In preparation).

To test this idea, we conducted a comparative analysis of trophic networks spanning diverse ecosystems. For each species, we quantified its meso-structural environment using a suite of local structural traits, including hierarchical asymmetry, weighted trophic position, and multiple forms of centrality (Allesina & Pascual, 2009; Brännström et al., 2011; Gravel et al., 2011). We then evaluated how these traits predict interaction outcomes and whether they reveal structural gradients consistent with selection. Finally, we compared meso-structural patterns with those obtained from aggregated networks to assess how analytical scale influences the detectability of structural signals.

Despite extensive work on global network structure, the scale at which selection acts within ecological networks remains unresolved. By focusing on meso-structural environments captured through egonets, we provide a framework that complements whole-network approaches and clarifies how local interaction geometry shapes selective pressures, revealing the structural domains where evolutionary signals become detectable.

## Methods

Formal framework: local structural selection in a directed trophic network We treat the food web as a directed, weighted graph.

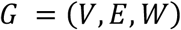

where V is the set of species, E is the set of predator–prey links (predator-prey), and W contains interaction weights.

To quantify structural selection at meso-structural scales, we analyze order-2 egonets, defined as induced subgraphs centered on a focal species and its neighbors. For species i, the order-2 egonet is:

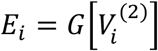

where 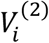 contains all nodes at distance ≤ 2 from i. Egonets capture local structural environments that are lost in aggregated networks. This framework parallels classical approaches to local selection analysis (e.g., Lande & Arnold, 1983) by quantifying how traits embedded in local structural geometry predict interaction outcomes.

The egonet based framework does not rely on trophic directionality; any network *G* = (*V, E*), including bipartite graphs, admits order-k egonets and local structural predictors. Thus, the approach generalizes directly to plant–pollinator, host–parasite, and other bipartite systems.

Local trophic metrics

For each egonet, we computed:

Weighted trophic level (wTL)

Using an iterative algorithm:

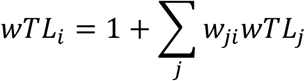

with guaranteed convergence via damping and tolerance thresholds.

Local hierarchy

Defined as the deviation of each species from the global hierarchical axis.

Structural predictors

For each ordered predator–prey pair (i, j) within an egonet we computed:

Δ hierarchy = hi - hj

Δ wTL = wTLi - wTLj

Degree product: kikj

Δ centrality (weighted betweenness)

Trophic similarity (Jaccard overlap of neighbors)

These quantities define local structural traits embedded in the geometry of the egonet, analogous to predictor variables in classical selection analyses.

Local structural models

For each egonet we evaluated two candidate mechanisms: M2: hierarchical–strength model

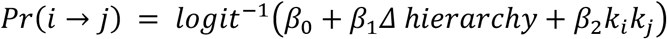

M4: hierarchical–motif model

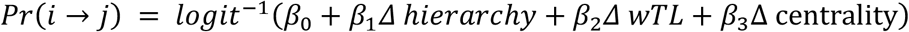

Models were fitted using binomial GLMs, and the best model for each egonet was selected using AIC. This approach parallels the logic of classic selection analyses (Lande & Arnold, 1983), where regression coefficients quantify the strength and direction of selection acting on traits—in our case, meso-structural traits.

## Extraction of structural signatures

For each species we extracted: the winning model (M2 or M4), the vector of standardized coefficients β, the set of local structural predictors. These β-vectors define local structural signatures, representing the selective landscape experienced by each species.

### Manhattan analysis

For each predictor we ranked species by β and plotted values across egonets. This identifies dominant mechanisms (predictors with long tails), structural outliers, and predictors with negligible influence, providing a structural analogue to classical selection-gradient visualization.

### PCA of structural signatures

We performed PCA on the matrix of standardized β-vectors to reveal dominant structural gradients, secondary axes of variation, and clustering of species by local structural environment.

### Data source

We used the trophic network available in the Web of Life repository (Bascompte et al., 2005), standardized to consistent species identifiers and imported as weighted directed graphs. For each network we extracted: number of species, number of links, connectance, and weighted adjacency matrix.

### Statistical Analysis

All analyses were conducted in R, version 4.5.1 (R Core Team, 2025). Data preprocessing consisted of standardizing interaction matrices, removing empty rows or columns, and normalizing interaction frequencies by total sampling effort when required. No data values were removed or modified beyond these transformations, and all preprocessing steps are fully documented in the accompanying code repository.

For each network, the units of analysis were: (i) individual interactions for aggregated networks, and (ii) meso structural domains for domain level analyses. Sample sizes (n) for each analysis correspond to the number of domains or interactions used and were reported in the figure legends.

Distributions of domain level metrics (e.g., domain size, internal density, structural deviation) were summarized using medians and interquartile ranges, as these variables were not normally distributed. For approximately symmetric distributions (e.g., aggregated connectance), means and standard deviations were reported. All measures of central tendency and dispersion are explicitly stated.

Comparisons between aggregated and domain level structural metrics were performed using non parametric tests (Wilcoxon rank sum or signed rank tests), given the non normality of the data. Two sided testing was used throughout. Effect sizes and 95% confidence intervals were computed using bootstrap resampling (10,000 iterations). No multiple testing corrections were required, as analyses were limited to a predefined set of structural metrics.

Assumptions underlying each statistical procedure (e.g., independence of domains, distributional properties) were evaluated using diagnostic plots and residual analyses. No missing data were present, and no imputation was performed.

All statistical procedures, including preprocessing, domain detection, structural selection quantification, and figure generation, are fully reproducible using the archived code and data.

### Software

All analyses were conducted in R using igraph (Csárdi & Nepusz, 2024), dplyr (Wickham et al., 2024), and custom scripts for similarity matrices, thresholding, null models, and selection analyses. All code will be archived in a public repository upon publication.

## Results

### Local structural environments reveal hidden heterogeneity

Although the aggregated representation of the network appeared structurally homogeneous, with weak gradients in global metrics such as degree or centrality (Fig. 1), order-2 egonets revealed substantial heterogeneity in local structural environments. Species differed markedly in the composition, asymmetry, and hierarchical arrangement of their immediate neighborhoods, even when these differences were not detectable at the whole-network scale. Definitions of all meso-structural traits are provided in Table S1. The construction of order-2 egonets and the workflow used to extract meso-structural traits are shown in Fig. 2. This pattern indicates that meso-structural analyses capture information that is not fully represented in global summaries, as reflected in the dispersion of egonet signatures along the first principal component (Fig. 3A).

**Fig. 1.**
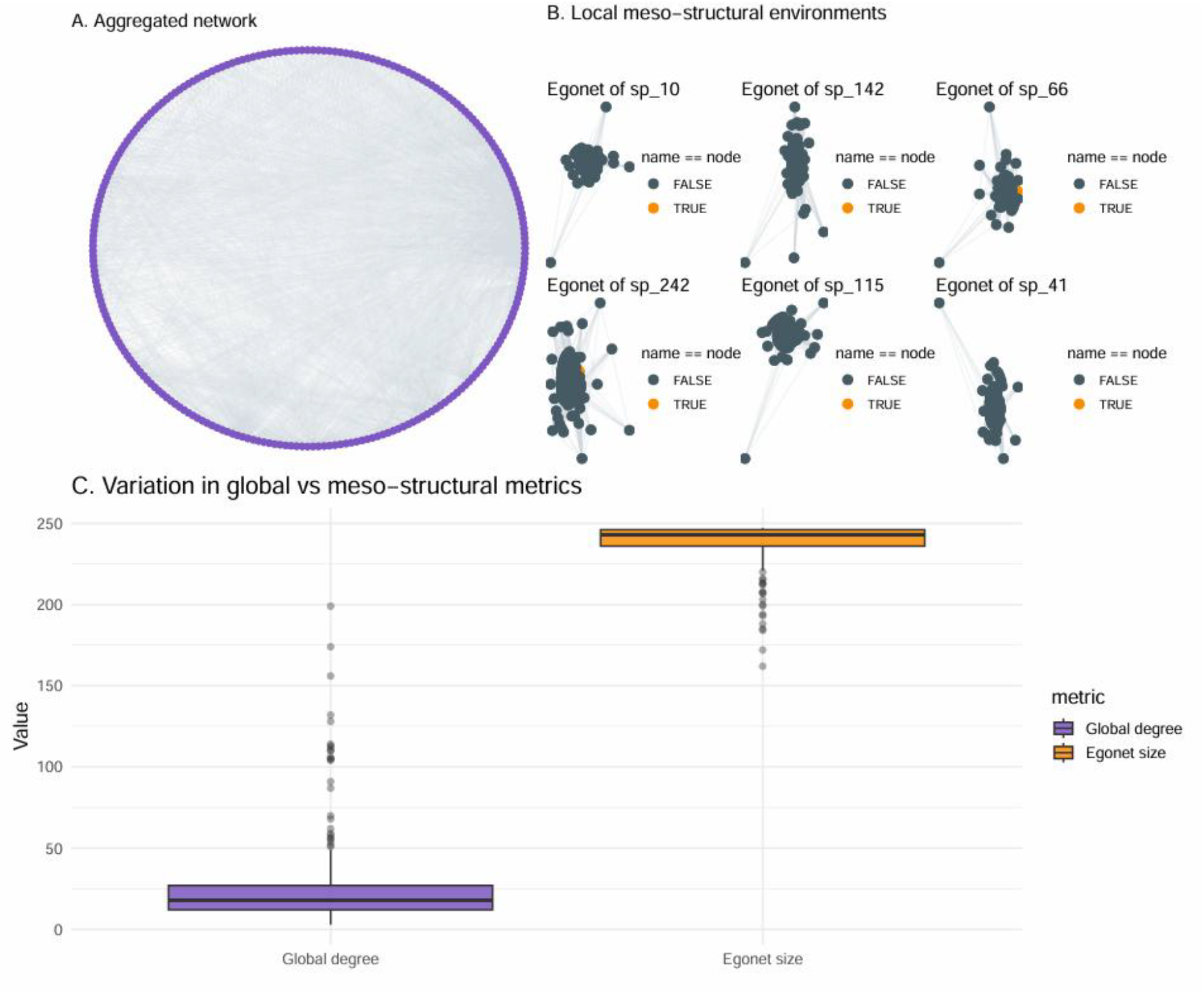
Meso-structural analysis reveals local structural environments hidden in aggregated networks. A) Aggregated trophic network showing a dense global topology with minimal apparent structural differentiation across species. B) Order-2 egonets for representative species illustrate the heterogeneity of local meso-structural environments, including variation in neighborhood size and connectivity patterns. C) Comparison between global degree and egonet size shows that meso-structural neighborhoods capture substantially greater variation than whole-network metrics, demonstrating that aggregated topology underestimates local structural complexity.

**Fig. 2.**
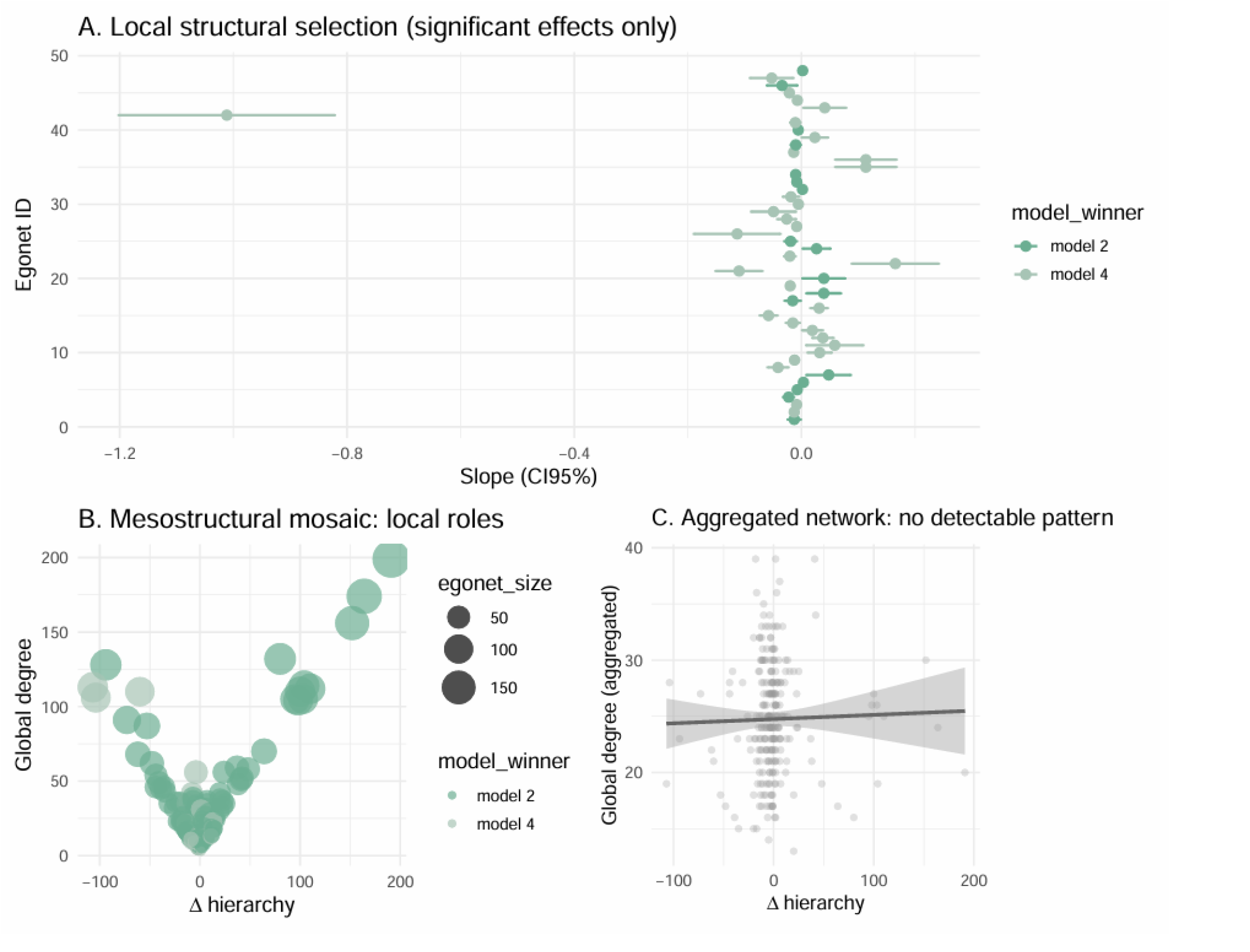
Local and meso-structural patterns reveal structural roles that disappear in the aggregated network. A) Local structural selection (significant effects only) showing slope estimates and 95% confidence intervals for individual egonets. Most effects cluster near zero, but a subset of egonets exhibit strong directional responses. B) Meso-structural mosaic of local roles, where Δ hierarchy and global degree define a V-shaped structural landscape. Egonet size increases toward the extremes of Δ hierarchy, indicating that meso-structural environments vary systematically across species. C) Aggregated network analysis shows no detectable relationship between Δ hierarchy and global degree, demonstrating that whole-network topology collapses the structural gradients evident at meso-structural scales.

**Fig. 3.**
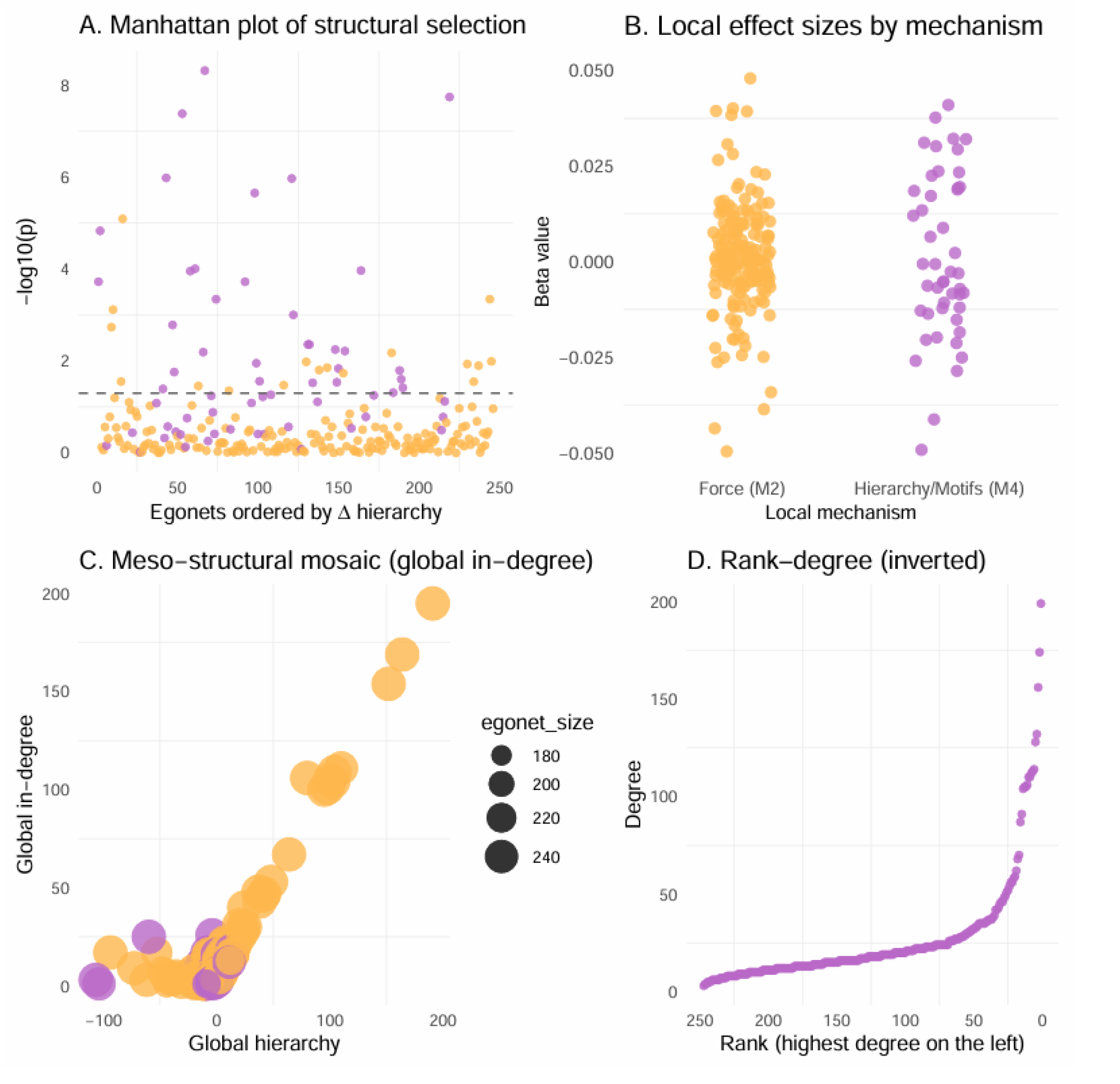
Meso-structural metrics reveal structural domains that retain selective gradients. A) Principal component analysis of meso-structural metrics shows two dominant axes of variation, with Δ hierarchy and egonet size contributing strongly to Dim1.B) K-means clustering identifies four meso-structural domains defined by combinations of Δ hierarchy and local degree, revealing a mosaic of distinct structural roles across species. C) Loadings on PC1 highlight Δ hierarchy as the primary driver of meso-structural differentiation, with local degree and egonet size providing secondary contributions. D) Global degree varies systematically across meso-structural domains, indicating that these domains retain structural gradients that are not detectable in the aggregated network.

### Hierarchical asymmetry dominates meso-structural variation

Among all meso-structural traits quantified, hierarchical asymmetry (Δ hierarchy) emerged as the dominant axis of variation (Fig. 3A). Species occupying asymmetric positions, where neighbors differed sharply in influence or connectivity, showed distinct local configurations relative to species embedded in more symmetric neighborhoods. Loadings on PC1 confirmed that Δ hierarchy contributed the strongest weight to the primary gradient (Fig. 3C; Table S2), whereas local degree, indegree, and out-degree contributed secondary structure. These results suggest that the relative arrangement of neighbors, rather than absolute global position, plays a central role in shaping local interaction geometry.

### Meso-structural domains reveal discrete local roles

Clustering of egonet signatures identified four discrete meso structural domains (Fig. 3B), each characterized by distinct combinations of hierarchical asymmetry and local connectivity. One domain contained highly asymmetric exporters, another concentrated asymmetric receivers, a third represented the neutral structural background of the network, and a fourth captured strongly hierarchical hubs. Extended trait distributions for each domain are shown in Table S3. These domains correspond to functional local roles that are not readily apparent in aggregated representations, where projection can smooth over underlying variation. Boxplots of global degree across domains further illustrate the retention of structural gradients within local environments (Fig. 3D).

### Structural selection is sparse but concentrated in hierarchical domains

Structural selection analyses revealed that most species were best explained by the neutral force model (M2; n = 186), exhibiting slopes near zero and high p-values, consistent with weak or absent directional structure (Fig. 4A). In contrast, a distinct subset of species aligned with the hierarchical/motif-based model (M4; n = 61), displaying more negative slopes and lower p-values (Fig. 4B). Although p-values were not uniformly low, model comparison showed that M4 provided a better explanation for these species, indicating that selective pressures are detectable but localized within the most asymmetric meso-structural domains. Table S4 includes all regression results and statistics related to model fit. The network shows near-neutral meso-structural dynamics, with only a few hierarchically embedded species showing structural selection.

**Fig. 4.**
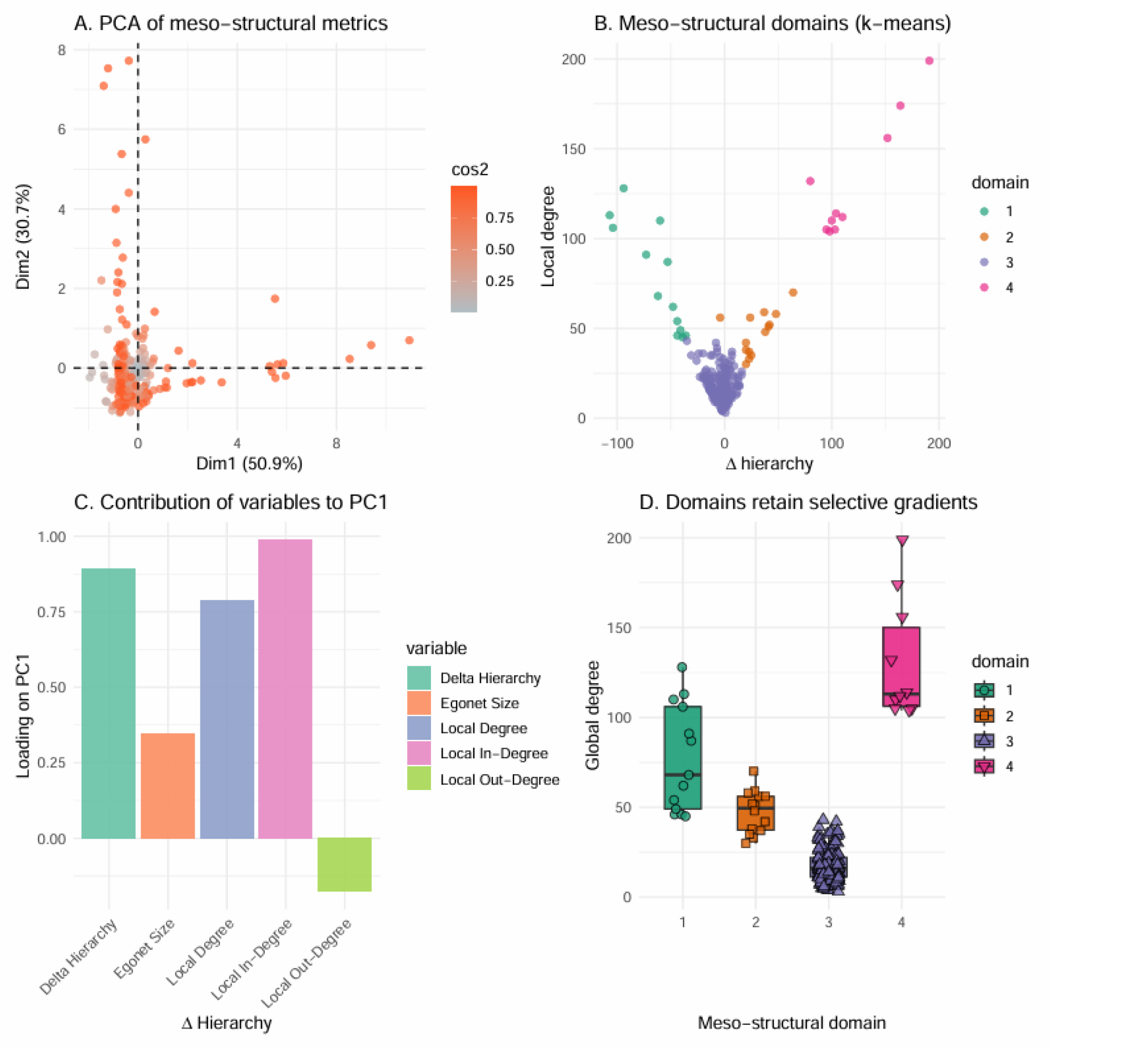
Structural selection emerges from hierarchical and meso-structural organization. A) Manhattan plot of structural selection showing ™log10(p) values for egonets ordered by Δ hierarchy. A subset of egonets exhibits significant structural effects, indicating localized selective pressures. B) Local effect sizes for two mechanisms—force-based (M2) and hierarchy/motif-based (M4)—demonstrate that hierarchical motifs generate stronger and more consistent directional effects. C) Meso-structural mosaic of global in-degree across the global hierarchy axis, with point size representing egonet size. Larger egonets cluster at hierarchical extremes, revealing systematic structural differentiation across species. D) Inverted rank–degree distribution showing that high-degree species occupy the upper hierarchical positions, reinforcing the link between global structure and meso-structural selective gradients.

Together, these results show that meso structure reveals hidden heterogeneity, organizes species into discrete local roles, and concentrates selective gradients in specific hierarchical domains. Analyses conducted at aggregated scales may overlook these patterns, not because selection is absent, but because it acts primarily at the scale of local interaction geometry rather than at the whole network level.

## Discussion

Our results reveal strong and consistent meso-structural differentiation across systems. Egonets exhibit clear axes of variation, dominated by hierarchical asymmetry, which predicts interaction probability and partner identity. In contrast, aggregated networks collapse these gradients, producing topologies that appear structurally uniform even when underlying meso-structural variation is substantial. This pattern helps clarify why previous analyses have struggled to detect structural selection: the relevant variation is not absent; it is simply masked when examined at aggregated scales.

By identifying meso-structure as the structural level at which selection operates, our study provides a mechanistic explanation for the limitations of whole-network analyses and establishes a framework for integrating structural selection into eco-evolutionary theory. Shifting the analytical focus to meso-structural scales reveals where selection acts, how structural constraints shape ecological opportunity, and why local interaction geometry, not global topology, determines the evolutionary consequences of species’ positions in trophic networks.

Our results demonstrate that structural selection emerges at meso-structural scales, where local interaction geometry retains the heterogeneity through which selective gradients can form (Bascompte & Jordano, 2007; Stouffer et al., 2012). This perspective helps resolve a long-standing puzzle in ecological network analysis: why whole-network metrics often fail to predict interaction outcomes or reveal consistent structural constraints. Aggregated networks collapse the variation that selection acts upon, producing topologies that appear homogeneous even when underlying meso-structural environments differ sharply among species (Pascual & Dunne, 2006; Poisot, 2016). This pattern also highlights a conceptual convergence between structural selection and classical natural selection: both processes filter variation through differential persistence, albeit acting on different substrates(Miranda-Pérez et al., In preparation).

Together, these results position structural selection as an evolutionary process acting on ecological structure, expanding how selection can be conceptualized in complex systems. Ego-centered networks, often referred to as “egonets” in modern network analysis, were originally introduced to characterize local neighborhoods (Wasserman & Faust, 1994); here we extend this concept to ecological networks formalizing meso-structural traits and evaluating whether selection acts specifically on variation at this level.

By identifying hierarchical asymmetry as the dominant axis of meso-structural differentiation, our study provides a mechanistic basis for understanding how local topology shapes ecological opportunity. Species embedded in asymmetric neighborhoods experience directional constraints on resource access, vulnerability, and partner identity, consistent with theoretical predictions that local structural imbalance shapes ecological dynamics (Allesina & Tang, 2012; Johnson et al., 2014). These constraints are difficult to detect in aggregated representations, where the relative arrangement of neighbors is lost. The prominence of hierarchical asymmetry across domains suggests that it reflects a general organizing principle of trophic structure rather than an idiosyncratic feature of particular food webs (Stouffer et al., 2005; Williams & Martinez, 2000). These insights have broad implications for eco-evolutionary theory. If selection acts primarily on meso-structural traits, then evolutionary responses should depend on the local configuration of interaction partners rather than on global network position. This view aligns with empirical evidence that species adapt to the immediate ecological context they encounter, rather than to the abstract topology of the entire community (Guimarães et al., 2017; Thompson, 2005). It also suggests that structural selection may operate even in systems where whole-network analyses detect no signal, simply because the relevant variation resides at a finer structural scale (Bartomeus et al., 2016; García*-*Callejas et al., 2018).

Our findings also clarify why classical network approaches have struggled to detect selective gradients. Metrics computed on aggregated networks implicitly assume that all species experience the same structural environment. This assumption is violated in real ecosystems, where interaction neighborhoods differ in composition, asymmetry, and hierarchical structure (Ings et al., 2009; Tylianakis et al., 2010). By focusing on meso-structure, we recover the gradients that drive interaction outcomes and reveal the structural heterogeneity that underlies ecological roles.

Future work should explore how meso-structural traits evolve, how they interact with species’ functional traits, and how they shape the assembly and stability of ecological communities. Integrating meso-structure into eco-evolutionary models may help illuminate how local constraints scale up to influence global patterns of diversity, resilience, and network architecture (Brännström et al., 2011; Gravel et al., 2011). More broadly, our framework provides a foundation for extending selection theory to structured ecological systems, where the geometry of interactions—not just their presence or absence—determines the evolutionary landscape. Although coevolution has been extensively documented in bipartite networks such as plant– pollinator and host–parasite systems (Bascompte & Jordano, 2007; Guimarães et al., 2017; Thompson, 2005), these studies typically focus on pairwise or guild-level interactions rather than on the structural environment of entire communities. Empirical work has shown that selective signals and structural heterogeneity are often detectable only at local or intermediate scales (García et al., 2024; Villa-Galaviz et al., 2012), supporting the idea that ecological interactions are shaped by the immediate configuration of partners rather than by global topology. This historical emphasis may reflect the fact that bipartite networks preserve more local heterogeneity than aggregated trophic webs, making selective gradients easier to detect (Ings et al., 2009; Tylianakis et al., 2010).

Our results show that this limitation is not inherent to trophic systems: when analyzed at the appropriate meso-structural scale, trophic networks also contain the structural variation upon which selection can act. This provides evidence that selection can operate on network-embedded properties at the community level, even when whole-network analyses detect no signal (Bartomeus et al., 2016; García*-*Callejas et al., 2018). By revealing that meso-structural traits carry selective gradients, our framework suggests that evolution in ecological networks is shaped not only by who interacts with whom, but by the geometry of the local neighborhoods in which those interactions occur. This insight motivates a broader synthesis in which eco-evolutionary dynamics emerge from the interplay between local structural constraints and global community architecture, encouraging a shift from aggregated representations toward meso-structural perspectives.

## Acknowledgments

We thank Jordi Bascompte for making his trophic network data publicly available and for an early conversation that inspired the initial direction of this work.

## Data, code, and materials availability

A permanent Zenodo repository will be created upon acceptance; the DOI will be added in the next version of the preprint.

## Author Contributions

Adán Miranda-Pérez designed the conceptual framework, implemented the analytical pipeline, conducted all computational analyses, and wrote the manuscript.

Citlalli H. Mendoza-Reyes contributed sustained conceptual support, provided intellectual input during the development of the framework, and supported the writing process through continuous discussion and critical engagement.

