## Supplemental Data 1 for "Structural selection is revealed only at meso‑structural scales"

Supplementary Material

Tables

Table S1. Summary of global and meso-structural metrics.

| Metric | Value |
|---|---|
| Number of nodes | 247 |
| Mean global degree | 26.73 |
| SD global degree | 28.76 |
| Mean egonet size | 237.94 |
| SD egonet size | 13.31 |
| Mean Δ hierarchy | 0 |
| SD Δ hierarchy | 31.67 |

† Global degree refers to the number of links per species in the aggregated network.
‡ Egonet size corresponds to the number of nodes in each order-2 egonet.
§ Δ hierarchy measures deviation from the global hierarchical axis.

Table S2. Correlations among meso-structural metrics.

| Metric 1 | Metric 2 | Correlation |
|---|---|---|
| Δ hierarchy | Local degree | 0.451 |
| Δ hierarchy | Local in-degree | 0.867 |
| Δ hierarchy | Local out-degree | −0.589 |
| Δ hierarchy | Egonet size | 0.182 |
| Local degree | Local in-degree | 0.836 |
| Local degree | Local out-degree | 0.456 |
| Local degree | Egonet size | 0.197 |
| Local in-degree | Local out-degree | −0.107 |
| Local in-degree | Egonet size | 0.222 |
| Local out-degree | Egonet size | −0.002 |

† Correlations computed using Pearson's r.
‡ Δ hierarchy measures deviation from the global hierarchical axis.
§ Egonet size corresponds to the number of nodes in each order-2 egonet.

Table S3. Mean meso-structural metrics across meso-structural domains.

| Domain | n | Mean Δ hierarchy | Mean local degree | Mean local in-degree | Mean local out-degree | Mean egonet size |
|---|---|---|---|---|---|---|
| 1 | 13 | −61.92 | 77.31 | 7.69 | 69.62 | 238.08 |
| 2 | 14 | 30.07 | 47.50 | 38.79 | 8.71 | 242.21 |
| 3 | 210 | −3.87 | 17.24 | 6.69 | 10.56 | 237.25 |

| | | | | | | |
|---|---|---|---|---|---|---|
| 4 | 10 | 119.70 | 131.10 | 125.40 | 5.70 | 246.30 |

† Domains correspond to k-means clusters identified in Fig. 3B.
‡ Δ hierarchy measures deviation from the global hierarchical axis.
§ Egonet size refers to the number of nodes in each order-2 egonet.

Table S4. Structural selection metrics across local mechanisms.

| Mechanism | n | Mean slope | SD slope | Mean p-value |
|---|---|---|---|---|
| Force (M2) | 186 | 0.00110 | 0.01366 | 0.54152 |
| Hierarchy/Motifs (M4) | 61 | −0.00681 | 0.13987 | 0.15741 |

† Slope values represent local structural selection coefficients estimated for each egonet.
‡ Mechanisms correspond to the two local models evaluated in Fig. 4B.
§ Mean p-values summarize the significance of structural effects across egonets.

Figures

Fig. S1. Distributions of meso-structural metrics used to characterize local structural environments.

A) Distribution of Δ hierarchy showing a sharp concentration around zero with extended tails, indicating heterogeneous hierarchical positioning across species. B) Egonet size distribution revealing substantial variation in the number of nodes within order-2 neighborhoods. C) Local degree distribution showing a right-skewed pattern, with most species having few local connections and a minority exhibiting high local connectivity. D) Local in-degree distribution highlighting asymmetries in incoming interactions within egonets. E) Local out-degree distribution showing that outgoing interactions are more constrained than incoming ones, with most species exhibiting low out-degree.

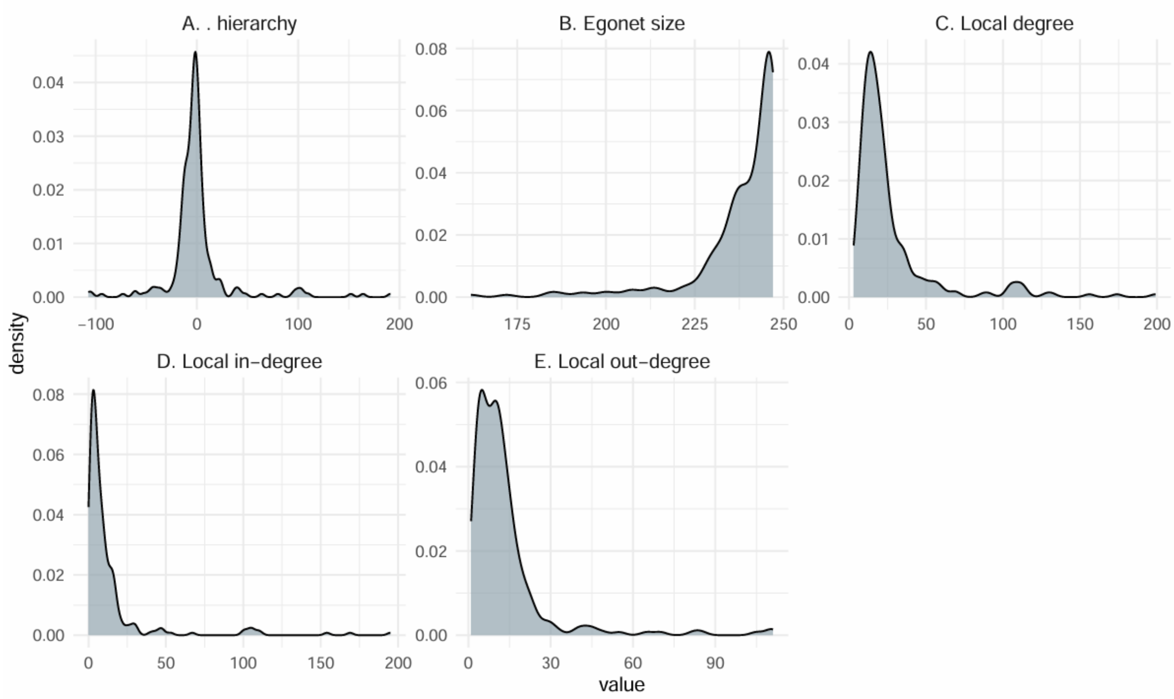
